# Arrhythpy: An Automated Tool to Quantify and Classify Arrhythmias in Ca^2+^ Transients of iPSC-Cardiomyocytes

**DOI:** 10.1101/2025.03.19.644093

**Authors:** Karim Ajmail, Charlotte Brand, Thomas Borchert, Sabine Rebs, Katrin Streckfuss-Bömeke

## Abstract

Arrhythmias constitute an intricate and clinically significant phenomenon of great importance in various research areas. Ca^2+^ homeostasis plays a pivotal role in forming rhythmic contractions in the heart, and its dysregulation has emerged as a critical component in developing arrhythmias. Until now, however, the quantification of arrhythmias has been limited to indirect measurements via Ca^2+^ sparks, electrophysiological parameters, or manual classification, which can lead to human bias. We aimed to develop an analysis platform that directly and automatically analyzes arrhythmias in human induced pluripotent stem cell-derived cardiomyocytes (iPSC-CMs).

Here, we present *Arrhythpy*, a robust and automated open-source program to quantify and classify confocal microscopy-based Fluo-4 Ca^2+^ transients to generate a measure of arrhythmia. In contrast to other automated and semi-automated analysis tools, that measure established parameters such as time-to-peak, *Arrhythpy* directly analyzes the degree of arrhythmia in a Ca^2+^ transient. We demonstrate its utility in monitoring Ca^2+^ transient-based arrhythmias in atrial and ventricular iPSC-CMs of healthy individual and cardiac disease patients, including dilated cardiomyopathy (DCM) and Takotsubo syndrome (TTS). *Arrhythpy* was validated by analyzing drug-treated iPSC-CMs, confirming the beating frequency effects of compounds that directly activate (Isoprenaline) or inhibit (Metoprolol) beta-adrenergic signaling. *Arrhythpy* analysis of iPSC-CMs of TTS patients recapitulated TTS phenotypes, including atrial arrhythmia that could be normalized with beta-blocker treatment. The program’s adaptable framework enables arrhythmic pattern analysis in various cell types using periodic dye-based line scan measurement techniques, applicable to single cells or layered cultures.

## I. INTRODUCTION

Arrhythmias, disturbances in the regular rhythm of the heart’s beating, represent a complex group of cardiac disorders that significantly impact the cardiovascular system. When the sinus rhythm is disrupted, the consequences can range from mild discomfort to life-threatening emergencies. Arrhythmias encompass a spectrum of irregularities, from slow (bradycardic) to fast (tachycardic) heartbeats, and originate in the atrium (supraventricular) or the ventricle (ventricular) (Masarone et al., 2017). They can be associated with various cardiac diseases as a primary manifestation or a secondary consequence of underlying cardiovascular conditions (Shah et al., 2005).

β1-adrenergic receptors are the most prevalent in the heart, comprising about 80% of cardiac β-adrenoreceptors. β1-receptors play a crucial role in regulating ventricular contractility and heart rate, exerting positive inotropic and chronotropic effects in response to catecholamines. However, these effects can also be arrhythmogenic. In contrast, beta-blockers counteract this by providing an antiarrhythmic effect, helping to stabilize heart rhythm. Metoprolol is a selective second-generation β1-blocker commonly used in clinical practice (Motiejunaite et al., 2021).

Dilated cardiomyopathy (DCM) is characterized by the enlargement and weakening of the heart’s chambers, particularly the left ventricle. This structural remodeling can have significant implications for the heart’s electrical system, making individuals with DCM susceptible to various arrhythmias (Sammani et al., 2020). While the Takotsubo syndrome (TTS) primarily affects the heart’s muscle contraction, it can also be associated with various arrhythmias, such as atrial fibrillation (El-Battrawy et al., 2021). Ca^2+^ plays a crucial role in regulating the heart’s electrical and mechanical activities, and disturbances in Ca^2+^ homeostasis can contribute to the development of arrhythmias. Abnormalities in Ca^2+^ release and reuptake in the sarcoplasmic reticulum (SR), as well as in key Ca^2+^-handling proteins, contribute to spontaneous Ca^2+^ release events and delayed repolarization, promoting the occurrence of an arrhythmogenic incidence (Ter Keurs & Boyden, 2007). Understanding the link between Ca^2+^ handling and the development of arrhythmias is crucial for unraveling the complexities of cardiac electrophysiology. Increased Ca^2+^ leakage from the SR can trigger a transient inward current, leading to arrhythmogenic delayed afterdepolarizations (DADs) and spontaneous proarrhythmic Ca^2+^ release. This process is further complicated by the late sodium current (INaL), which facilitates Na^+^ influx and increases Ca^2+^ influx through the Na^+^-Ca^2+^-exchanger (NCX). This elevation in cytosolic Ca^2+^ concentration enhances ryanodine receptor 2 (RyR2) Ca^2+^ release events, contributing to the arrhythmogenic potential (Hartmann et al., 2023; Hartmann et al., 2024).

The advent of human induced pluripotent stem cell (iPSC) technology has enabled researchers to reprogram somatic cells into a pluripotent state on a patient-specific level and, therefore, to investigate the pathophysiology of various, predominantly genetic disorders. The iPSC-based cardiac disease models offer insights into disease mechanisms, paving the way for developing targeted and personalized treatment strategies (Karakikes et al., 2015). A specialized tool to analyze abnormal Ca^2+^ homeostasis in iPSC-CMs would enable researchers to quantify arrhythmias.

To this end, we developed *Arrhythpy*, a user-friendly automated open-source pipeline for analyzing Ca^2+^ transients of iPSC-CMs implemented in the open-source programming language Python (G Van Rossum, 1995). It provides a nuanced characterization of arrhythmic behavior and assigns a numerical value to each transient as a measure of arrhythmia. *Arrhythpy* can also distinguish between tachycardic and bradycardic beats and automatically classifies transients based on the quantification of bradycardia and tachycardia. The User Interface and source code can be found at https://github.com/Karim-Ajmail/Arrhythpy.

## II. RESULTS

### Using *Arrhythpy* to analyze and quantify arrhythmias of atrial and ventricular iPSC-CMs

To demonstrate the capabilities of *Arrhythpy*, we examined line scans obtained via Fluo-4 Ca^2+^ measurements of iPSC-CMs of an individual described as healthy (CTRL) (Borchert et al., 2017), a patient with RBM20 mutation-based DCM (Rebs et al., 2020) and a patient with TTS (Borchert et al., 2017).

According to previously published protocols, we generated ventricular (for CTRL and DCM) (Bengel et al., 2021) and atrial (for TTS) (Hartmann et al., 2023) iPSC-CMs and submitted them to confocal Fluo-4 measurements to acquire Ca^2+^ transients as sketched in Fig. 1A.

**Fig. 1:**
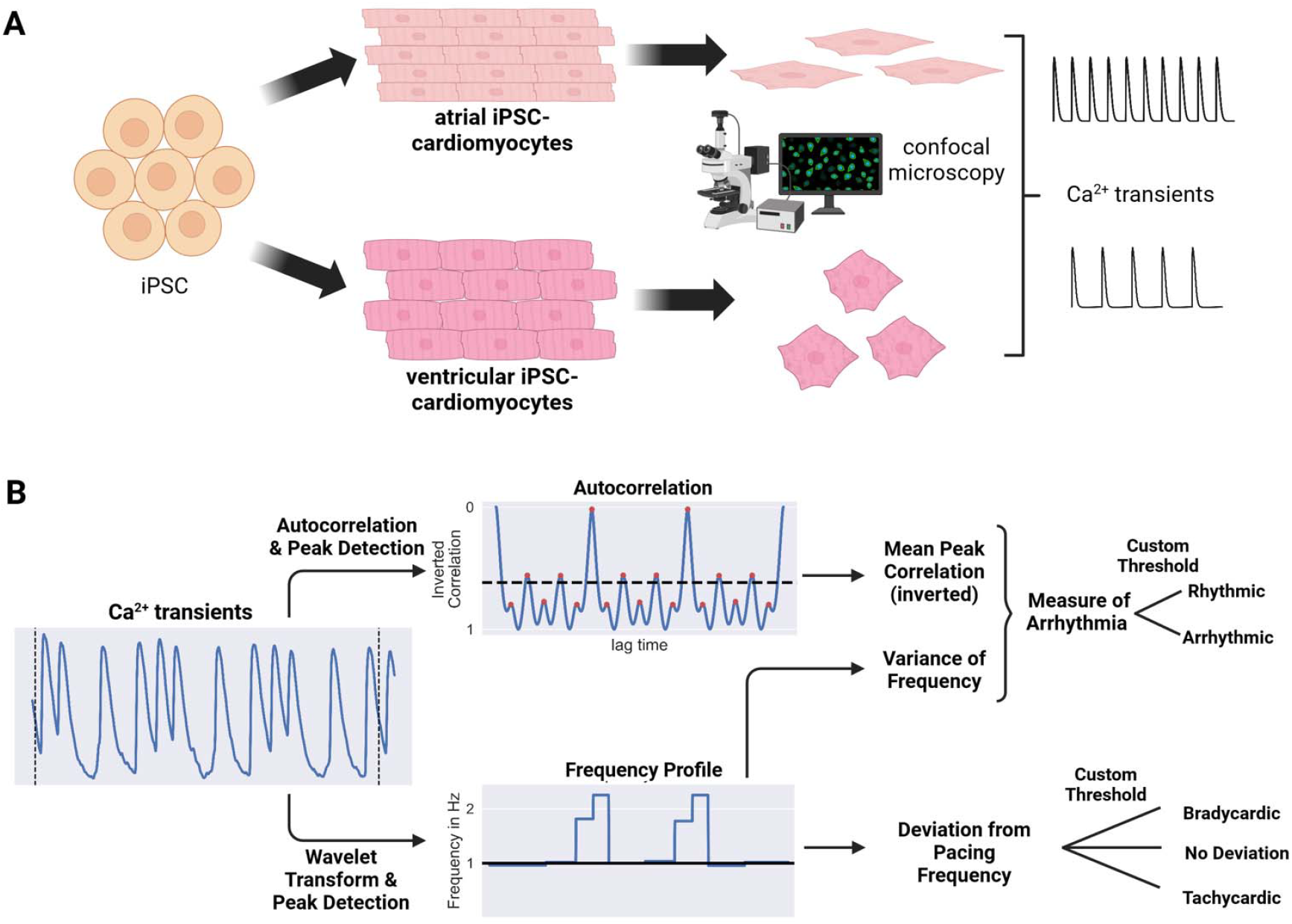
Working principle of *Arrhythpy*. A) Overview of differentiation into atrial or ventricular iPSC-CMs and subsequent preparation for Fluo-4 Ca^2+^ measurement. B) Working principle of *Arrhythpy*. Ca^2+^ transients are obtained as depicted in A). The line scan is first cropped to the first and last root to create a periodic boundary (dashed line). For capturing variations in frequency and deviation from the pacing frequency, a wavelet transformation combined with peak detection and interpolation is performed resulting in the frequency profile. Secondly, the autocorrelation of the transient is recorded to probe the periodicity of the signal. To this end, all peaks of the autocorrelation are detected and averaged. The measure of arrhythmia is obtained with the frequency variance obtained via the wavelet transformation. For both methods, the user defines a custom threshold for the classification into a total of six classes (namely Rhythmic, Arrhythmic, Tachycardic Rhythmic, Tachycardic Arrhythmic, Bradycardic Rhythmic, and Bradycardic Arrhythmic).

In order to quantify arrhythmias in transients, we first defined arrhythmias in a computer-readable manner by incorporating two aspects of arrhythmias. Firstly, arrhythmias mainly cause temporal fluctuations in frequency within a transient. Secondly, variations in the shape of each peak of a transient, regardless of the frequency, are a strong indicator of arrhythmias. To consider these two aspects, *Arrhythpy* consists of two parts, depicted in Fig. 1B. The first part obtains frequency information via wavelet transformation (lower) and computes a frequency profile, which gives the frequency at any time point of the transient. Standard algorithms for peak detection to determine the width of each peak are often only used on preselected rhythmic transients and fail at strongly arrhythmic transients. The wavelet transformation provides a more robust method for determining the widths and frequencies within one transient. The second part (upper) computes a measure of the periodicity of the shape of each peak using the autocorrelation function. Averaging each peak of the autocorrelation function and inverting the scale yields the mean peak correlation. The lower the mean peak correlation, the more similar the peaks of the transient. The deviation from the pacing frequency of the experiment is used as a measure of tachycardia and bradycardia. By combining the variance of the frequency profile and a measure of periodicity obtained by autocorrelation, we find a robust method to assign a numerical value (Measure of Arrhythmia) that describes how arrhythmic a transient is. Additionally, setting an empirical threshold on the measure of arrhythmia enables automatic classification into arrhythmic and rhythmic. Similarly, setting a threshold on the deviation from the pacing frequency allows us to distinguish between tachycardic beating, bradycardic beating, or neither. In total, *Arrhythpy* is capable of classifying each transient into six categories, namely rhythmic, arrhythmic, tachycardic rhythmic, tachycardic arrhythmic, bradycardic rhythmic, and bradycardic arrhythmic. Fig. S1 shows an exemplary transient as well as frequency profile and inverted autocorrelation for each class. Table S1 lists the corresponding parameters obtained by analyzing the transients with *Arrhythpy*. We also provided a tool to conveniently determine the optimal thresholds and assess the accuracy of *Arrhythpy*.

To validate the successful differentiation into subtype-specific iPSC-CMs, we performed immunostainings, flow cytometry measurements, and quantification of atrial and ventricular-specific markers 60 days post-differentiation according to established protocols (Bengel et al., 2021), (Pabel et al., 2022), (Hartmann et al., 2023). For both atrial and ventricular iPSC-CMs, we found typical striated sarcomere structures in immunofluorescence staining of α-actinin and titin (Fig. 2A, 2B). Flow cytometry for cardiac Troponin T (cTNT) found on average 85.15% atrial iPSC-CMs and 91.77% ventricular iPSC-CMs (Fig. 2C) positive for cTNT. To further demonstrate the specific differentiation into atrial and ventricular cells, respectively, we analyzed cells positive for the atrial marker myosin light chain atrium (MLC2a), and for the ventricular marker myosin light chain ventricle (MLC2v) (Fig. 2D, E). 89,25% of atrial cells were positive for MLC2a and negative for MLC2v (MLC2a+, MLC2v-), whereas 64.49 % of ventricular cells were negative for MLC2a and positive for MLC2v (MLC2a-, MLC2v+) (Fig. 2F). Expression of sub-type specific markers (atrial: NR2F2; ventricular: MLC2v) confirmed their successful differentiation in atrial and ventricular iPSC-CMs (Fig. 2G).

**Fig. 2:**
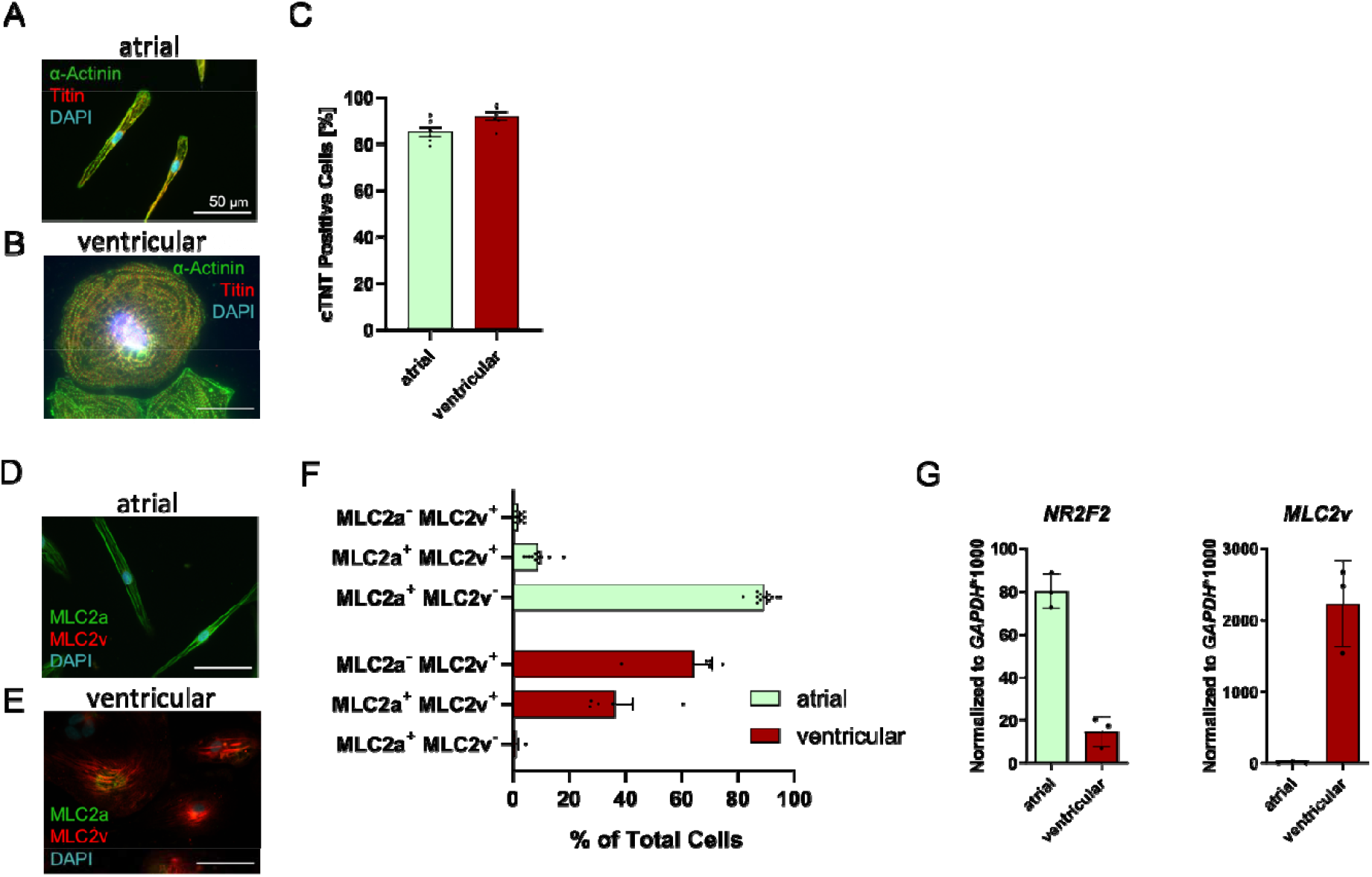
Quality control of differentiation into atrial and ventricular iPSC-CMs. A) and B) Representative immunostainings of α-actinin (green), titin (red) and 4’,6-Diamidino-2-phenylindole (DAPI) (blue) of atrial (A) and ventricular (B) iPSC-CMs. Atrial iPSC-CMs display the characteristic elongated morphology, whereas ventricular iPSC-CMs assume a rounder morphology (scale bar: 50 µm). C) Flow cytometry data of cTNT confirming the differentiation into both atrial and ventricular iPSC-CMs. D) and E) Representative immunostainings of MLC2a (green), MLC2v (red) and DAPI (blue) of atrial (D) and ventricular (E) iPSC-CMs (scale bar: 50 µm). F) The majority of atrial cells (89.25%) were MLC2a+ and MLC2v-, with 1.9% being MLC2a- and MLC2v+. In contrast, ventricular cardiomyocytes showed 1.86% as MLC2a+ and MLC2v-, and 64.49% as MLC2a- and MLC2v+. (G) qPCR-based quality control for atrial marker nuclear receptor subfamily 2 group F member 2 (NR2F2) and ventricular marker MLC2v.

### Quantification of the effect of Isoprenaline on arrhythmia development in ventricular iPSC-CMs

To verify the applicability of *Arrhythpy*, we first investigated the influence of the exogenous catecholamine isoprenaline (Iso) on arrhythmogenesis via β1-adrenoceptor activation. Diastolic Ca^2+^ leaks from the SR, also described in the literature as Ca^2+^ sparks, are a known proarrhythmic trigger (Dridi et al., 2020). Ca^2+^ sparks are visible in line scans as puncta marking leakage from the SR into the cytosol, indicating increased diastolic Ca^2+^ leakage into the cytoplasm. Fig. 3A and B show that increasing Iso concentrations lead to an increased frequency of sparks in healthy ventricular iPSC-CMs, as illustrated by the exemplary images of line scans in Fig. 3A (marked by white arrows) and the quantification of sparks in Fig. 3B. Averaging these line scans yields the transients of paced iPSC-CMs used for the analysis by *Arrhythpy* (Fig. 3C). Fig. 3D-I present the six parameters that *Arrhythpy* provides as analysis output to analyze whether *Arrhythpy* can reliably characterize changes in iPSC-CMs under Iso-stimulation.

As expected, increasing the concentration of Iso to 1 µM leads to a rise in the mean frequency (Fig. 3D). However, saturating the Iso concentration at an unphysiological concentration of 5 mM does not result in a further increase in mean frequency, unlike observed for sparks (Fig. 3B). Fig. 3D and E show that most deviations from the pacing frequency are due to tachycardia, with the strongest effect at 1 µM Iso. At the same time, bradycardia plays a minor role, even though significant differences between high and low Iso concentrations were observed. The frequency variance (Fig. 3G) and mean peak correlation (Fig. 3H) highlight different aspects of arrhythmias. The frequency variance captures strong tachycardic events occurring at 1 µM, whereas the mean peak correlation characterizes differences in the shape of the peaks. At high, cytotoxic and unphysiological concentrations of Iso (5 mM), the cells cannot beat fast enough, leading to slight irregularities in the peaks (as shown in Fig. 3C at 5 mM). These irregularities are detected by the mean peak correlation. Finally, combining the frequency variation and the mean peak correlation yields the measure of arrhythmia (Fig. 3I), indicating that up to 100 nM Iso, no change in arrhythmia as characterized by *Arrhythpy* is induced, but above 1 µM Iso, arrhythmias arise.

### Analysis of arrhythmias in ventricular iPSC-CMs of an RBM 20 - DCM patient

To investigate commonalities and differences in arrhythmogenesis between healthy control and DCM patients (Rebs et al., 2020), we analyzed and quantified arrhythmias in ventricular iPSC-CMs of an RBM20 mutation (p.R634W)-based DCM patient under basal conditions as well as after Iso-treatment (1 *µ* M). RBM20 mutations are known to cause highly penetrant and arrhythmogenic cardiomyopathies with mainly non-sustained ventricular tachycardia and atrial fibrillation as an accompanying trait (Jordà et al., 2021).

We compared the effect of Iso on DCM iPSC-CMs in comparison to control iPSC-CMs (Fig. 4). Exemplary line scans and transients for DCM in basal and Iso-treated conditions can be found in Table S2. As anticipated, catecholamine stress induced by Iso-treatment (1 µM) resulted in significantly increased tachycardic events in both healthy and DCM iPSC-CMs. We observed a significant increase in mean frequency (Fig. 4A) and tachycardic beating (Fig. 4B) after Iso-treatment. In contrast, there was no significant increase in the fraction of bradycardic beating cells after Iso stress in DCM cells (Fig. 4C). The variance of the beating frequency indicates a significantly higher degree of arrhythmia after Iso-treatment in DCM iPSC-CMs compared to basal conditions (Fig. 4D), whereas the mean peak correlation shows a trend of a higher degree after Iso-treatment without significance (Fig. 4E). Naturally, this leads to a significant increase in the combined measure of arrhythmia after Iso-treatment for both, DCM and CTRL (Fig. 4F). However, even under basal conditions, DCM iPSC-CMs display an arrhythmic behavior, as can be seen by the increase in tachycardia, frequency variance, mean peak correlation, and measure of arrhythmia when compared to CTRL iPSC-CMs under basal conditions (Fig. 4B, D, E, F). Interestingly, Iso-treatment results in a slightly different arrhythmia phenotype between DCM and CTRL. For control cells, Iso-treatment leads to clear additional tachycardic beating events, which is reflected by the increased mean frequency, tachycardia as well as frequency variance. However, DCM iPSC-CMs display a significantly higher mean peak correlation upon Iso-treatment than the control. This phenotype of DCM iPSC-CMs stimulated at 1*µ*M is similar to the control cells stimulated at 5mM Iso (Fig. 3), which indicates a hypersensitivity of DCM iPSCs-CMs to Iso-stimulation.

**Fig. 3:**
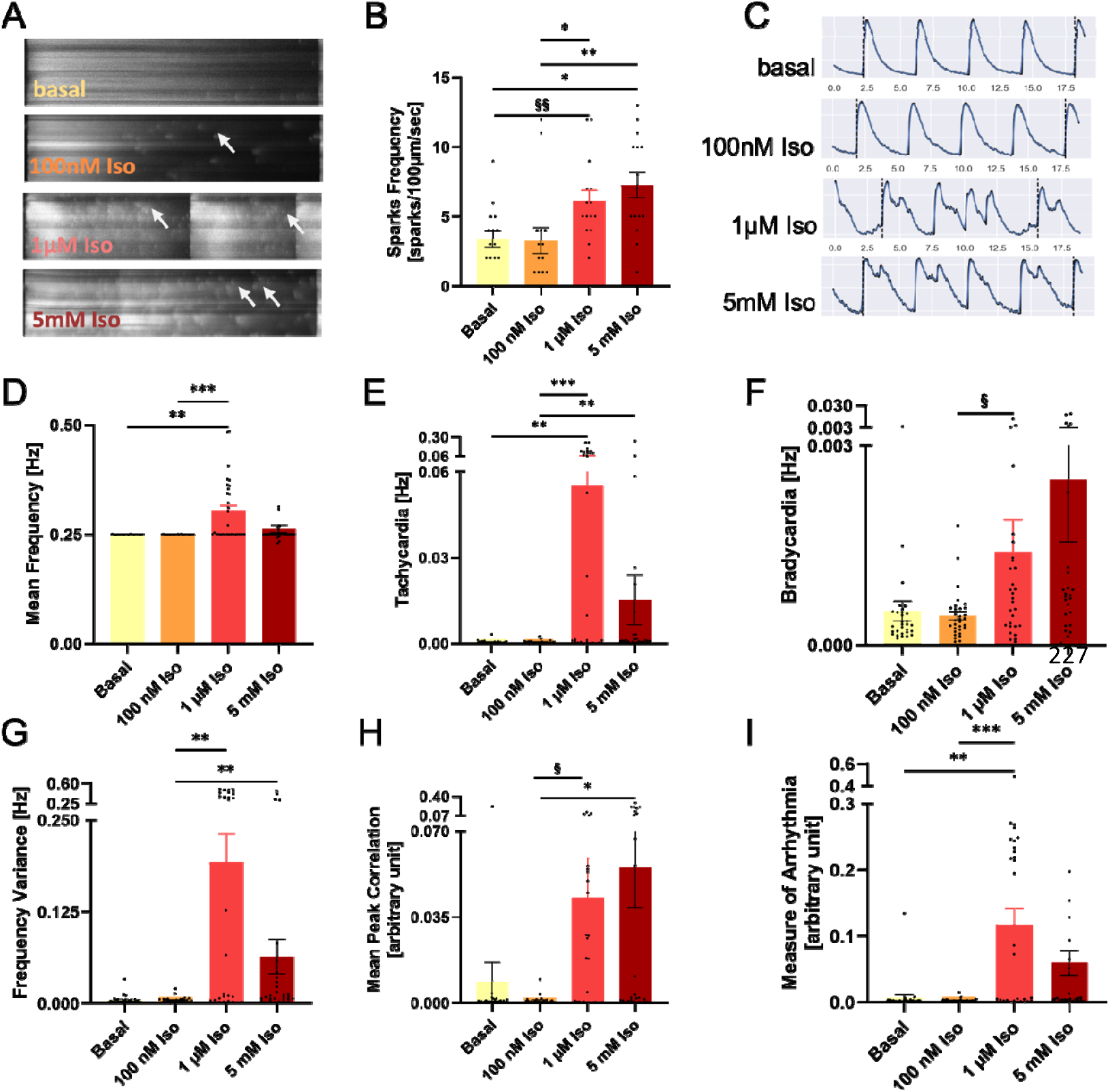
Iso-induced changes in Ca^2+^ dynamics and arrhythmogenic activity in ventricular iPSC-CMs. A) Original representative line scans of control ventricular iPSC-CMs illustrating a spontaneous arrhythmogenic diastolic Ca^2+^ wave (sparks), marked by white arrows (line scan duration: 1s). B) Above an Iso concentration of 100nM spark frequency increases with increasing Iso concentrations. C) Ca^2+^ transients obtained by averaging the respective line scans shown in A, for *Arrhythpy* analysis. D-I: Parameters obtained by *Arrhythpy* script. D) Mean frequency, obtained by averaging the frequency profile, peaks at 1 µM Iso and decreases to near baseline at an unphysiological high concentration of 5 mM Iso. E) & F) Positive (tachycardic) and negative (bradycardic) deviations from the pacing frequency. Stimulation of iPSC-CMs with Iso concentrations higher than 100 nM leads to dominant tachycardic deviation, indicating that most cells exhibit tachycardic behavior rather than bradycardia. G) Frequency variance. High variance at 1 µM Iso indicates clear beating events, that deviate from the pacing frequency, which decreases at a cytotoxic concentration (5 mM). H) Mean peak correlation, measuring dissimilarities between peaks in Ca^2+^ transients, peaks at 5 mM (unphysiological control), indicating varying transient peak shapes. I) Measure of arrhythmia computed from G) and H). Data were compared using non-parametric ANOVA (Kruskal-Wallis) for multiple comparisons (*) and unpaired non-parametric t-test (Mann-Whitney) (§) to calculate p-values.

**Fig. 4:**
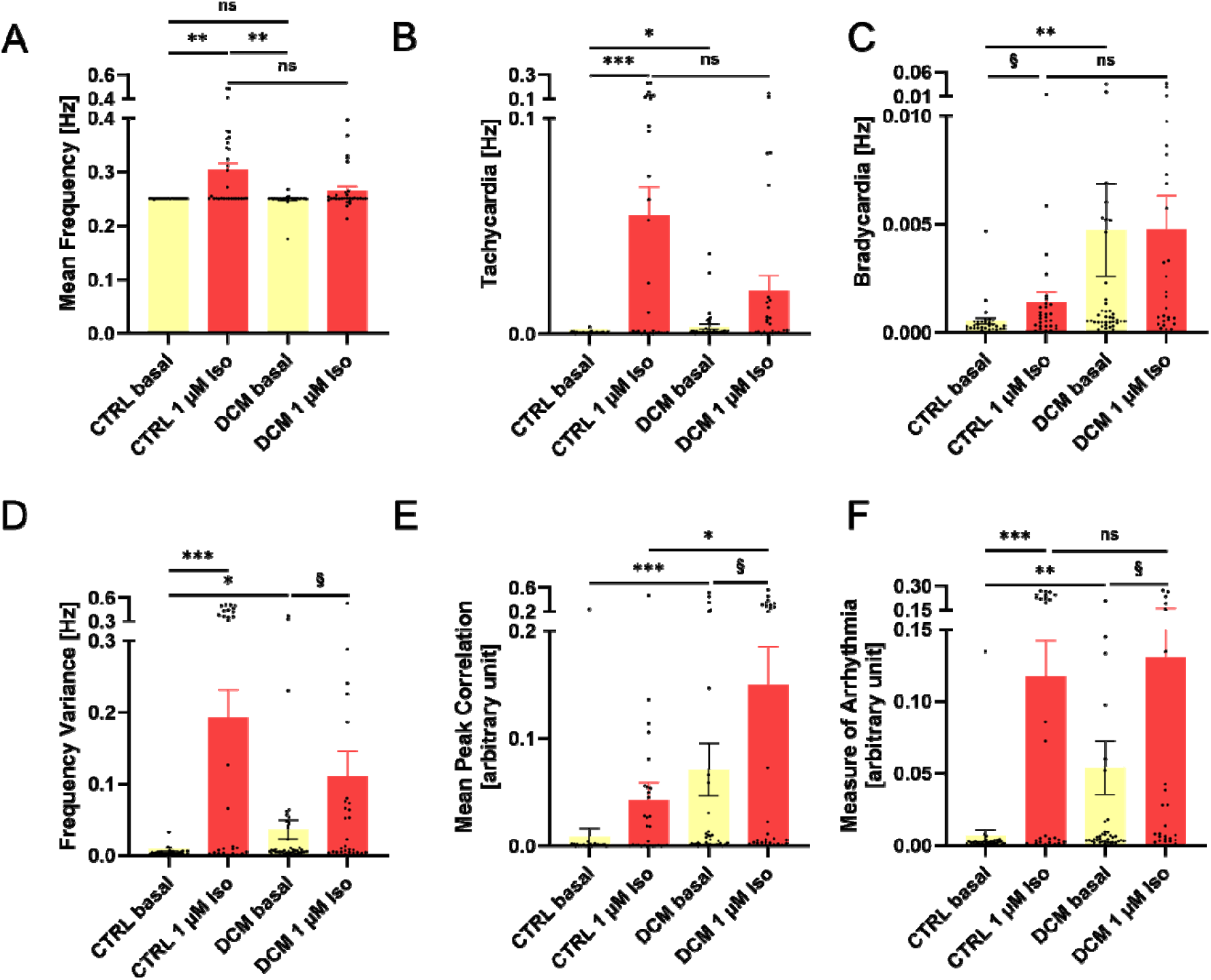
Iso-induced effects on frequency, tachycardia, and arrhythmia metrics in DCM iPSC-CMs. A) Mean frequency, B) Tachycardia, C) Bradycardia, D) Frequency Variance and E) The Mean Peak Correlation of CTRL and DCM iPSC-CMs without and with Iso-treatment (1 µM). F) Measure of arrhythmia computed from D) and E). Iso: 1 µM. Data were compared using non-parametric ANOVA (Kruskal-Wallis) for multiple comparisons (*) and unpaired non-parametric t-test (Mann-Whitney) (§) to calculate p-values.

### High arrhythmias in atrial hiPSC-CMs of a Takotsubo patient

Further, we tested the capability of *Arrhythpy* to correctly analyze and classify line scans of atrial cardiomyocytes paced at a different frequency.

Atrial fibrillation occurs in 18% of TTS cases and persists throughout both the acute phase and the recovery period (El-Battrawy et al., 2017). To explore whether *Arrhythpy* can detect this form of arrhythmia, we generated atrial iPSC-CMs from a TTS patient (Borchert et al., 2017) and conducted confocal Ca^2+^ measurements. iPSC-CMs were treated with Iso to induce catecholaminergic stress and subsequently relieve the arrhythmia with the beta-blocker Metoprolol. Exemplary line scans and their corresponding transients of atrial health and TTS iPSC-CMs are depicted in Table S2. These line scans were subsequently analyzed using *Arrhythpy* to assess the presence of arrhythmias.

In our atrial iPSC-CM TTS model, we quantitatively compared arrhythmias (Fig. 5) under basal conditions, after Iso-treatment (50 nM) and after subsequent treatment with the beta-blocker Metoprolol (5 µM), which is clinically used as an antiarrhythmic drug. Catecholamine stress via Iso-application increases the mean beating frequency (Fig. 5A) and a deviation from the pacing frequency towards a tachycardic rhythm (Fig. 5B) in atrial TTS iPSC-CMs. We did not detect a shift towards a bradycardic beating after Iso-treatment (Fig. 5C). Both, the variance of the beating frequency (Fig. 5D) and the mean peak correlation (Fig. 5E) indicate a higher degree of arrhythmia after Iso-treatment compared to basal conditions in atrial TTS cells, resulting in a high measure of arrhythmia for TTS cells compared to CTRL cells (Fig. 5F). To test whether *Arrhythpy* is capable of detecting a change in arrhythmic beating after application of antiarrhythmic drugs, we subsequently treated the Iso-stressed cells with Metoprolol. In all parameters computed by *Arrhythpy*, the basal phenotype was restored, confirming the efficacy of Metoprolol’s antiarrhythmic effect. However, even under basal conditions, atrial TTS iPSC-CMs display an arrhythmic behavior, as can be seen by the increase in bradycardia, mean peak correlation, and measure of arrhythmia when compared to atrial CTRL iPSC-CMs under basal conditions (Fig. 5C, E, F).

**Fig. 5:**
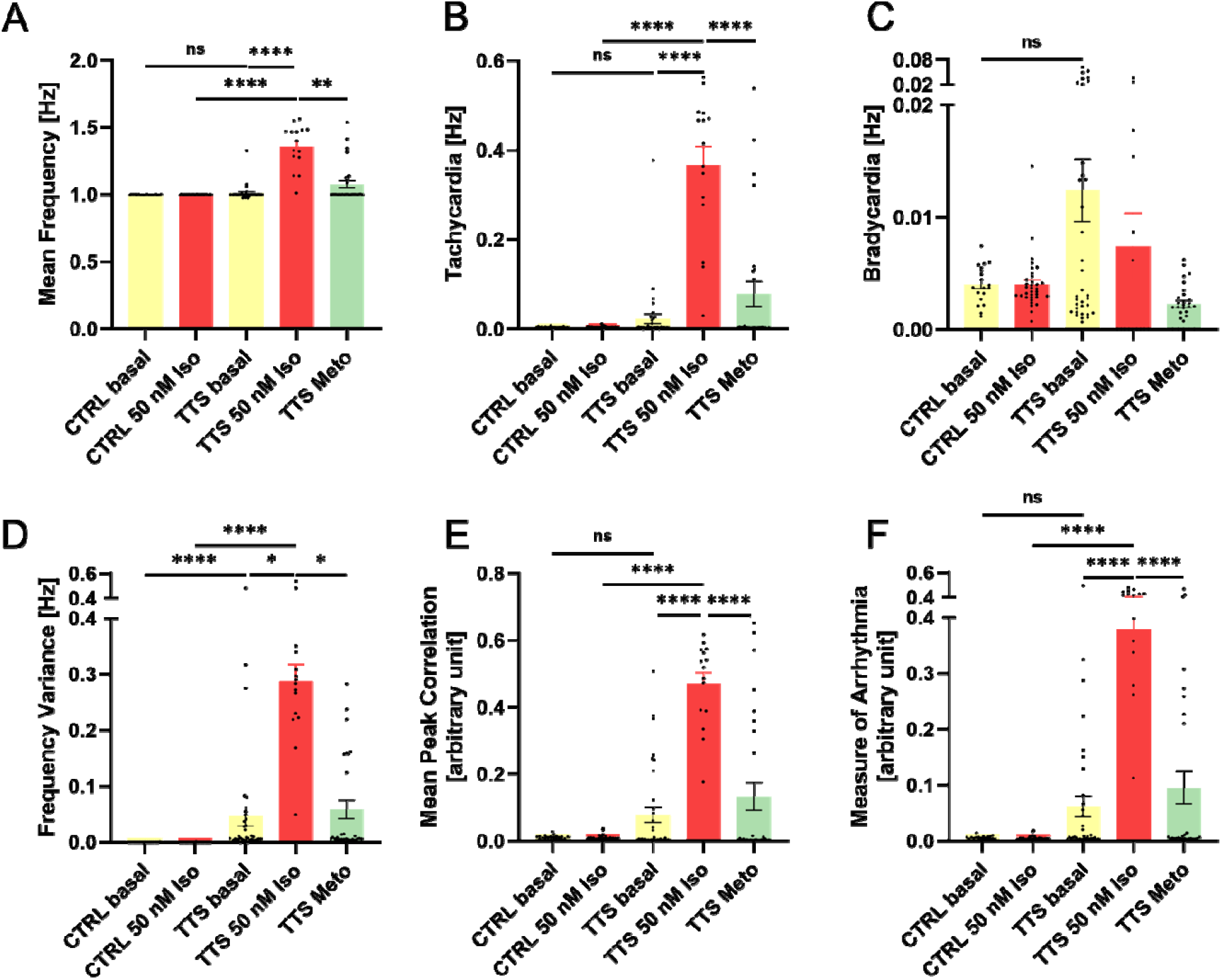
Arrhythmia assessment in atrial TTS iPSC-CMs. A) The Mean Frequency; B) & C) the positive and negative deviation from the pacing frequency, respectively. D) Variance of the frequency profile. E) Mean Peak Correlation and F) Measure of Arrhythmia computed from D) and E) all confirm the strong trend of an increase in arrhythmia upon Iso-stimulation and subsequent rescue to basal levels by Metoprolol treatment in TTS iPSC-CMs. Iso: 50nM; Metoprolol: 5µM. Data were compared using non-parametric ANOVA (Kruskal-Wallis) for multiple comparisons (*) and unpaired non-parametric t-test (Mann-Whitney) (§) to calculate p-values.

### Automated classification of arrhythmias in iPSC-CMs

The parameters calculated by *Arrhythpy* can further be used to classify transients into six classes by thresholding the measure of arrhythmia, separating into arrhythmic and rhythmic, as well as the deviation from the pacing frequency, separating into tachycardic, bradycardic, and paced. Note, that in this context, rhythmic describes a regular periodic transient regardless of its frequency. For example, a transient beating regularly at 2 Hz despite being paced at 1 Hz is classified as tachycardic rhythmic, a transient that changes frequency between 0.5 Hz and 0.75 Hz would be bradycardic arrhythmic and a transient whose frequency fluctuates around the pacing frequency is classified as arrhythmic. Fig. 6 shows the classified transients obtained from the datasets presented in this work (ventricular CTRL: Fig. 6A; ventricular DCM: Fig. 6B; atrial TTS: Fig. 6C). The classification highlights the effect of Iso, as we can see the clear shift from rhythmic towards tachycardic arrhythmic beating in CTRL, DCM and TTS cells. In the control cells 1 µM promotes tachycardic arrhythmias and only a small fraction of arrhythmic transients, whereas at unphysiological Iso concentrations (5 mM) the number of arrhythmic cells increased. In contrast, almost half of DCM iPSC-CMs treated with 1 µM Iso are arrhythmic. Similarly, TTS iPSC-CMs show a very strong shift toward tachycardic arrhythmic behavior, which can be rescued to basal levels upon treatment with Metoprolol.

**Fig. 6:**
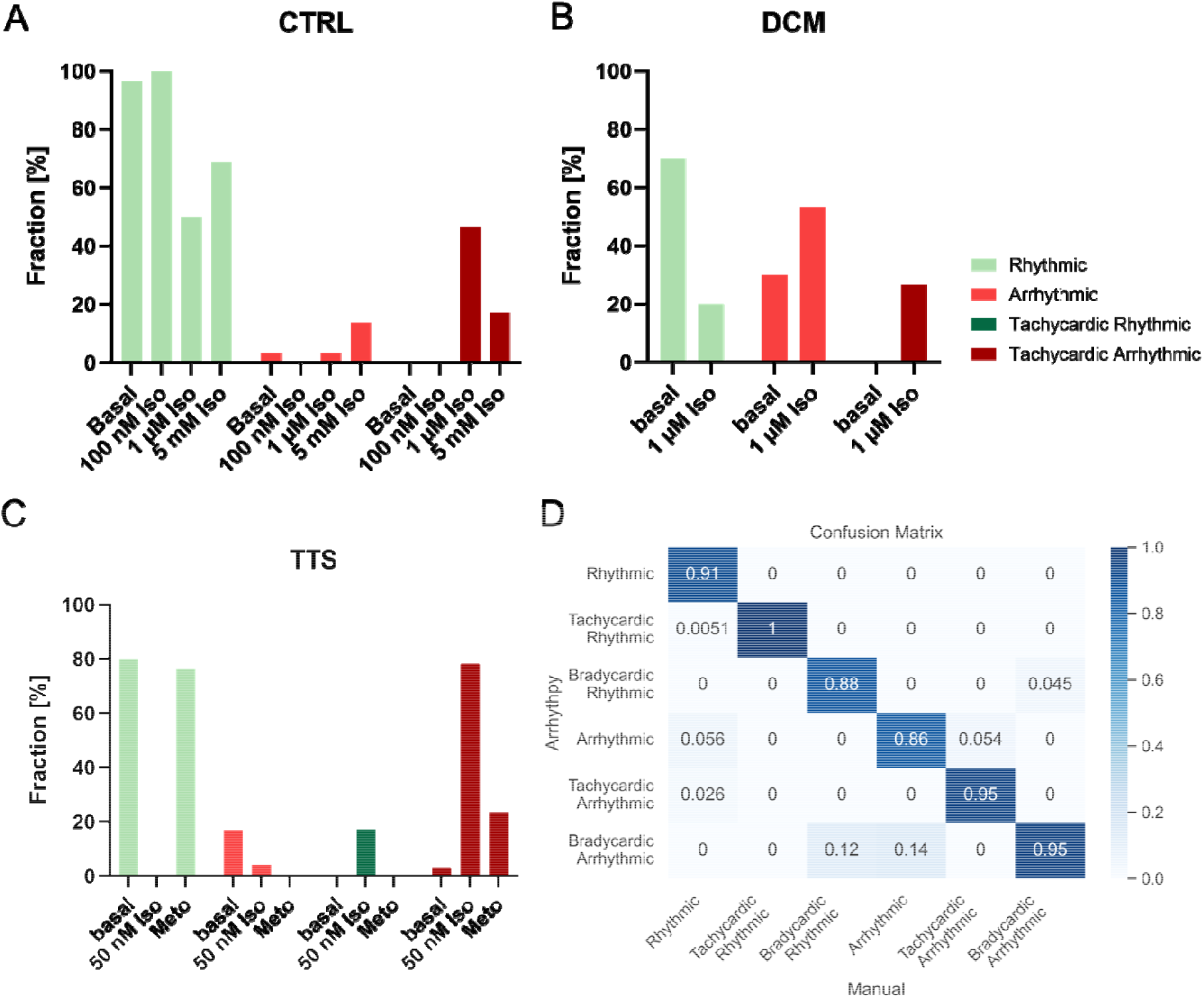
Threshold-based classification of iPSC-CMs Ca2+ transients. Iso increases the fraction of tachycardic arrhythmic beating significantly for all, ventricular CTRL (A), ventricular DCM (B), and atrial TTS (C) iPSC-CMs. Metoprolol rescues this to basal levels for TTS. D) The confusion matrix shows the relative misclassification for each class compared to a manual classification. The dominant diagonal indicates a good performance of *Arrhythpy*.

The classification also allows for assessing the accuracy of *Arrhythpy* by manually classifying transients and comparing the results to the prediction of *Arrhythpy*. By adjusting the two thresholds for classification, we achieved an overall accuracy of 92.4% in total. The confusion matrix, shown in Fig. 6D confirms the absence of any systematic misclassification. In contrast, most machine learning-based methods depend on a large quantity of consistently labelled data. However, in a laboratory setting, the amount of labeled data is limited, and most transients are often rhythmic, leading to strongly imbalanced data. To test this, we trained a simple convolutional neural network (CNN) on a small manually labeled dataset to classify the transients in the six categories that *Arrhythpy* also provides. Fig. S2 compares the confusion matrices of *Arrhythpy* (Fig. 6D, Fig S2A) and the CNN classification (Fig. S2B). It is clear, that the CNN favors rhythmic classification due to the imbalanced nature of the dataset. Further, since bradycardic rhythmic and tachycardic rhythmic are the rarest classes, CNN does not learn to distinguish between bradycardic/ tachycardic rhythmic and bradycardic/tachycardic arrhythmic. This results in a biased classification, highlighting the advantage of *Arrhythpy* over machine learning approaches in the case of an insufficiently balanced dataset.

## III. DISCUSSION

In this paper, we demonstrate that *Arrhythpy*, an open-source, user-friendly Python-based program, is capable of directly quantifying arrhythmias in Ca^2+^ transients, a task that was previously only possible with indirect measures such as Ca^2+^ spark analysis.

Here, we demonstrated the use of *Arrhythpy* in electrically paced ventricular and atrial iPSC-CMs, that were imaged in line scan-mode on a confocal microscope system with the Fluo-4 Ca^2+^-dye. However, since *Arrhythy* does not discriminate between different dyes or microscope settings, the script can be used on any Ca^2+^ investigation set-up which includes different cell types, different cell densities (monolayer or single-cell), different Ca^2+^-dyes (Fluo-4, ratiometric dyes: Fura-2 or Indo-1), genetically encoded Ca^2+^ sensors and different pacing set-ups (electrically or optically). There are only two requirements for *Arrhythpy*: Firstly, the required input for the script are either images (.lsm or .tiff) or raw data (.csv) of Ca^2+^ transients. Secondly, the cells need to be paced since *Arrhythpy* correlates an expected pace with the measured Ca^2+^ transients. Notably, the use of fluorometric dyes like Fluo-4 is advantageous compared to ratiometric dyes because imaging kinetics are faster and, in combination with a confocal linescan, allow spatial-temporal detection of highly localized Ca^2+^ events such as Ca^2+^ sparks (Guatimosim et al., 2011).

Many automated and semi-automated analysis tools have been used to characterize Ca^2+^ transients (Psaras et al., 2021), (Oguntuyo et al., 2022). However, although the common parameters obtained, such as time-to-peak, RT50, etc., often are correlated to an arrhythmic phenotype, they do not directly characterize arrhythmia. *Arrhythpy* provides a **direct measure** for arrhythmia. Further, since these tools are based on peak-detection algorithms, they work best on rhythmic transients and are not robust against strong variations in the transients. Therefore, strongly arrhythmic transients are often not used for kinetics analysis. With this preselection, valuable biological information might be lost and biased towards a more rhythmic phenotype. *Arrhythpy* is independent of the kinetics analysis provided by many of these tools and hence covers additional information about the arrhythmic phenotype of the cells.

Data-driven approaches have come to dominate the automated analysis of arrhythmias, especially in ECGs (Ebrahimi et al., 2020), due to their undeniable success in cases where there is a large amount of data available. In a laboratory setting, the amount of consistently labeled data on Ca^2+^ transients is very limited and often includes mostly rhythmic transients, leading to a drastic decrease in performance. *Arrhythpy* does not require training as it is based on first principles. Further, most data-driven approaches only provide a classification, whereas *Arrhythpy* provides an array of parameters for quantitative assessment and classification.

We investigated whether *Arrhythpy* can quantify and classify arrhythmias by analyzing line scans obtained in Fluo-4 Ca^2+^ measurements of iPSC-CMs under different conditions, treatments, and genetic disease backgrounds with an externally applied pace of 0.25 Hz for ventricular and 1.0 Hz for atrial iPSC-CMs. We first set out to quantify the effect of Isoprenaline (Iso) on arrhythmias since Iso is known to increase arrhythmic behavior via β1-adrenoceptor activation (Novak et al., 2015). Indeed, the primary response at 1 µM Iso is a sharp increase in beating frequency and a higher number of tachycardic beating cells in ventricular CTRL iPSC-CMs. The beating frequency decreases at an unphysiological Iso concentration of 5 mM, while the shape deviation within a transient remains high, indicating irregularities. As Iso concentrations increase, cytotoxic effects also a r i s e. At 5 mM Iso (unphysiological control), the iPSC-CMs show the highest levels of diastolic Ca^2+^-leaks without a corresponding rise in measurable arrhythmia. The elevated diastolic Ca^2+^ levels might overwhelm the cells’ capacity for detectable beating, leading to irregular transient morphology. This finding supports integrating two distinct aspects of arrhythmias into a combined measure to effectively capture both irregularities. The unexpected observation of slightly increased bradycardia in Iso-stressed cells can be explained by a somewhat longer pause following several tachycardic events.

In the ventricular iPSC-CM model, DCM cells exhibit more arrhythmic behavior than control cells under basal conditions, as shown by increased tachycardia, frequency variance, mean peak correlation, and overall arrhythmia. Iso-treatment, similar to CTRL iPSC-CMs, induced tachycardic beating with an increased mean frequency and variance in beating frequency, emphasizing the universal response to catecholamine stress. Two important conclusions can be drawn from the data on DCM. Firstly, the basal levels of arrhythmic events are higher in the DCM model compared to the control. Secondly, the fingerprint of the parameters generated by *Arrhythpy* in the DCM cells is comparable to control cells stimulated with 5mM Iso. This leads to the conclusion that DCM entails higher sensitivity to Iso. The results reinforce the potential of *Arrhythpy* as a versatile tool for quantifying arrhythmias in various cardiac settings.

In the atrial TTS iPSC-CM model, we paced cardiomyocytes at 1.0 Hz. We conducted measurements under basal conditions, after Iso-treatment, known to induce catecholamine stress, and after treatment with the antiarrhythmic drug Metoprolol. After the application of Iso, we observed a significant increase in the mean beating frequency, indicative of a tachycardic rhythm. Notably, the absence of a shift towards bradycardic beating after Iso-treatment in contrast to the DCM cells, underscores the specific response of the TTS cells. Our quantitative analysis demonstrated an increased variance in beating frequency and an altered mean peak correlation, collectively highlighting a higher degree of arrhythmia post Iso-treatment. The subsequent administration of the antiarrhythmic drug Metoprolol led to a significant reduction in the number of arrhythmic cells. The combined measure of arrhythmia, incorporating frequency variance and mean peak correlation, provided a comprehensive evaluation of arrhythmic events. Furthermore, we found that the basal levels of arrhythmic events are higher in the TTS model than the control, primarily based on bradycardic behavior as known from the left ventricular ballooning during a TTS event.

## Conclusion

In conclusion, *Arrhythpy* enables automated quantification and classification of arrhythmias in iPSC-derived cardiomyocytes. The observed trends in ventricular CTRL and DCM, as well as in atrial TTS models, highlight the significance of *Arrhythpy* in elucidating the complex dynamics of arrhythmias in different anatomical regions and disease states of the heart. Therefore, we offer a valuable platform for analyzing, quantifying, and classifying the occurrence of arrhythmias in paced Fluo-4-obtained Ca^2+^ line scans of patient-specific iPSC-CM models. Although the platform is optimized for Ca^2+^ transients of iPSC-CMs, it can be readily adapted to other cell types and experimental modes. With modern cardiovascular research transitioning from fundamental mechanistic discoveries to therapeutic applications, we expect that novel software such as *Arrhythpy* will further accelerate translational cardiovascular medicine.

## IV. METHODS

### A. Cell Culture

Human induced pluripotent stem cells (iPSCs) from a healthy individual (Borchert et al., 2017, JACC), a TTS patient (Borchert et al., 2017), and a DCM patient (Rebs et al., 2020) were used for cardiac differentiation.

The iPSCs were cultured at 37 °C with 5% CO2 in Essential8 (E8) Medium (Thermo Fisher Scientific) on Geltrex-coated (Thermo Fisher Scientific) 6-well plates (CytoOne). Once the cells reached 90% confluency, they were passaged in ratios ranging from 1:8 to 1:14 by incubating with Versene (Thermo Fisher Scientific) for 3–5 minutes, followed by resuspension in E8 Medium supplemented with 2 µM Thiazovivin (TZV) (Merck Millipore) and transferred to freshly coated plates. For cardiac differentiation, iPSCs were seeded on 12-well plates (CytoOne) at a density of 30,000 cells per well. Upon reaching 80-90% confluency, differentiation was initiated using Cardio Differentiation medium (RPMI 1640 Medium + GlutaMAX, 0.05% human recombinant Albumin, 0.02% L-ascorbic Acid 2-Phosphate) supplemented with 4 µM CHIRR99021 (Merck Millipore) to inhibit GSK3 (day 0). After 48 hours, the medium was replaced with Cardio Differentiation medium containing 5 mM inhibitor of Wnt production-2 (IWP2) (Merck Millipore) (Lian et al., 2013). For ventricular differentiation, IWP2 supplemented medium was incubated for 48 h as previously described (Borchert et al., 2017). For atrial differentiation, 1 µM retinoic acid (Sigma-Aldrich) was added on days 3, 4 and 5 as previously described (Cyganek et al., 2018). Cells were changed in Cardio Differentiation medium for two days and then refreshed every other day with Cardio Culture medium ((RPMI 1640 Medium supplemented with 2% B27 serum-free supplement 8 Thermo Fisher Scientific)) until spontaneous beating was observed. The cells were then replated onto 6-well plates at a density of 600,000-800,000 cells per well. For digestion, the cells were treated with 0.25% Trypsin-EDTA (Thermo Fisher Scientific) for five minutes, neutralized with inactivated fetal calf serum (FCS) (Sigma-Aldrich), centrifuged at 200 g for five minutes, and resuspended in Cardio Digestion medium (80% Cardio Culture medium, 20% inactivated FCS, 2 µM TZV). Following established protocols, the cells underwent a selection process using glucose-deprived Cardio Selection medium (RPMI 1640 Medium, 0.05% human recombinant Albumin, 0.02% L-ascorbic Acid 2-Phosphate, 4 mM Lactate/HEPES) for 3-5 days, with medium changes every other day until pure cardiac morphology was observed (Tohyama et al., 2013). Post-selection, the cells were cultured in Cardio Culture medium for 60 days to enhance maturation before measurements. In minimum, three cardiac differentiations were used for each analysis.

### B. Flow Cytometry

The efficiency of cardiac differentiation was verified by flow cytometry for cardiac troponin T (cTNT). Successful differentiation was defined by more than 85% cTNT-positive cells. At day 60, the cardiomyocyte monolayer was dissociated into single cells using 0.25% Trypsin-EDTA for 10 minutes at 37 °C, then transferred to a BD Falcon flow tube containing FCS. Following centrifugation at 200 g for five minutes and washing the cells twice with phosphate-buffered saline (1xPBS) (Thermo Fisher Scientific), cells were fixed with 4% Paraformaldehyde (ROTH) at room temperature for 20 minutes. After another centrifugation step and PBS wash, samples were stored at 4 °C in blocking solution (1% BSA in 1xPBS) (Thermo Fisher Scientific). Cells were divided into blank, secondary antibody control, and cTNT sample groups. Cells were resuspended in FACS buffer (1% BSA and 0.1% Triton X-100 (Sigma-Aldrich) in PBS). The cTNT sample was treated with the primary antibody (cTNT, 1:500, monoclonal mouse IgG (Thermo Fisher Scientific)) in FACS buffer and incubated overnight at 4 °C. The following day, cTNT and control samples were washed three times with FACS buffer and incubated with the secondary antibody (Alexa Fluor 488 anti-mouse, 1:1000, donkey anti-mouse IgG (Life Technologies)) for 45 minutes at room temperature in the dark. Samples were washed with 0.2% BSA/PBS and resuspended in PBS for analysis on a BD Biosciences FACS Canto II (settings: 10,000 events, Forward Scatter 228V, Side Scatter 440V, and 488 Alexa Fluor Laser 390V). The threshold for cTNT-positive cells was determined using the blank sample and the secondary antibody-only control was subtracted from the cTNT results.

### C. Immunostaining

For each differentiated cell line, immunostainings for MLC2v (1:200, polyclonal rabbit, (Proteintech)), MLC2a (1:200, monoclonal mouse IgG2B, (Synaptic Systems)) α-Actinin (1:500, Monoclonal mouse IgG1, (Sigma-Aldrich)), and Titin M-line M8/M9 (1:750, polyclonal rabbit IgG, (Myomedix)) were conducted to confirm the successful differentiation. To this end, 20,000 cardiomyocytes were plated on Geltrex-coated cover slips (R. Langenbrinck). After one week, they were fixed with 4% Paraformaldehyde for immunostaining and stored overnight at 4 °C in blocking solution (1% BSA in 1xPBS) (Thermo Fisher Scientific). The cover slips were then incubated with the primary antibody in a blocking solution containing 0.1% Triton X-100 and left overnight at 4 °C. Subsequently, the cells were treated with the secondary antibody for one hour, followed by 10-minute incubation with 4’,6-Diamidino-2-phenylindole (DAPI) (Sigma-Aldrich). Finally, the cover slips were mounted onto microscope slides using Fluoromount-G (Southern Biotech) and imaged using a Zeiss Axio Observer.Z1.

### D. Ca^2+^ Measurements

To assess Ca^2+^ fluctuations, iPSC-CMs underwent an initial loading with Fluo-4 am (2.5 µM, Invitrogen) supplemented with Pluronic F-127 at 0.05% w/v (Thermo Fisher Scientific) for 30 minutes, followed by 15 min incubation in Tyrode’s solution (5.4 mM KCl, 140 mM NaCl, 1 mM MgCl_2_, 10 mM HEPES, 10 mM glucose, 1.8 mM CaCl_2_, pH 7.4). Utilizing a LSM 720 confocal microscope (Zeiss) equipped with a Myopacer device (IonOptix), measurements were conducted with cells paced at 18 V for 3 ms at 1 Hz for atrial cells and 0.25 Hz for ventricular cells. The following parameters, as per the software Zen 2009, were employed: argon ion laser excitation at 488 nm with 1.0 – 2.2%, 700 gain and offset 0. For the line scan function, an area in the cytoplasm of a cell was chosen, and the line scan mode was configured to 20.000 cycles for ventricular cells and 10.000 cycles for atrial cells, 12 bits unidirectional, zoom factor 3, 512 pixels, no delay, and maximum scan speed. The images were analyzed using the Fiji software and sparks were analyzed using the ImageJ Plugin Sparkmaster.

### E. Details of *Arrhythpy*

#### 1. Wavelet transformation to obtain time resolved frequency information

Frequency information of a signal is commonly assessed by the Fourier transformation (FT), where the time series signal is decomposed into a sum of plane waves. The coefficient or the weight of the corresponding frequency mirrors the prevalence of this frequency in the signal. However, FT can only assess global frequency information and fails to accurately decompose non-stationary frequency information i.e. transients, that contain changing frequencies. These transients are of great interest, since they represent arrhythmias. Therefore, we use the wavelet transformation (WT), which compromises frequency resolution with additional time information. Hence, the output of the WT is two dimensional; one axis is frequency and one axis is time. This 2D plot is called coefficient map, whereas FT only results in a 1D coefficient signal. In more detail, WT is based on localized waves called wavelets. A plethora of wavelets is available, but for our purpose the Morlet wavelet empirically proved to be the most suitable choice. The wavelet depends on two variables, the shift *a* representing the time axis and the scale *b* representing the frequency axis. Hence, the convolution of the wavelet with the signal is computed for various scales. The maxima and minima in the coefficient map represent the dominant frequency component at that time point, where maxima represent the concave part of the signal and minima the convex part. Therefore, *Arrhythpy* assigns each maximum to its corresponding minimum to obtain an estimate of the width of an entire peak by averaging the scale of the minimum and the scale of the maximum. Note, that due to the nature of wavelets, the extrema have to be weighted in the average so that the length of the signal is equal to the sum of all widths. The widths or scales can readily be converted to frequencies and are interpolated with the nearest neighbor method. Using this method, one can obtain a time resolved frequency profile as depicted in Fig 1B. The positive and negative deviations from a given pacing or eigen frequency are used as a measure for tachycardia and bradycardia, respectively. Additionally, the variance of frequencies with the frequency profile (VF) is part of the measure of arrhythmia.

#### 2. Autocorrelation to obtain the measure of periodicity

However, the obtained frequency information does not contain information about the periodicity of the signal, since a signal, that is classified as rhythmic according to section IV E 1 might fluctuate around the pacing frequency. Therefore, an additional measure is needed to discriminate between rhythmic and arrhythmic signals regardless of the actual frequency. In order to develop a shape independent measure for periodicity, we calculate the autocorrelation of the signal (AC). For a perfectly periodic signal the normalized autocorrelation peaks to one regardless of shape or frequency, whereas non-periodic signals result in a decreased correlation. To avoid boundary artefacts, we implement a periodic boundary by appending the signal to itself at the first and the last root of the signal. The peaks of the autocorrelation, as shown in Fig. 1B, are detected, averaged and inverted to obtain a numerical value between 0 and 1 (mean peak correlation), with 0 representing perfect periodicity and 1 a completely random number.

As mentioned above, to capture different aspects of arrhythmias, the two measures VF and mean peak correlation are combined into a single value between 0 and 1. Applying a custom threshold to the FD and the AM yields 6 categories, that a signal can be classified into. Although the thresholds are set empirically, we also provide an optional tool to determine the best thresholds based on manually classified transients.

### F. Statistical Analysis

Statistical analysis was conducted using the analysis function in GraphPad Prism9. Differences between two independent groups were evaluated with an unpaired non-parametric t-test (Mann-Whitney). For comparisons involving more than two independent groups, a non-parametric one-way analysis of variance (ANOVA) (Kruskal-Wallis) was applied, followed by Dunn’s post hoc test to adjust for multiple comparisons. Statistical significances are indicated by * p<0.05, ** p<0.01, *** p<0.001 and ^****^ p<0.0001.

## Author Contributions

Conceptualization, K.A. and C.B.; Data curation, K.A. and C.B.; Formal analysis, K.A. and C.B., T.B. and S.R.; Funding acquisition, K.S.-B; Investigation, K.A. and C.B.; Methodology, K.A. and C.B T.B., S.R., and K.S.-B.; Project administration, K.S.B.; Resources, K.S.B.; Software, K.A. and C.B.; Supervision, K.S.-B.; Validation, K.A. and C.B, S.R., T.B., and K.S.-B; Visualization, K.A. and C.B.; Writing-original draft preparation, K.A., C.B., and K.S.-B. All authors have read and agreed to the published version of the manuscript.

## Informed Consent Statement

Informed consent was obtained from all subjects involved in the study.

### Acknowledgments

We thank Yvonne Metz and Johanna Heine for their excellent technical assistance.

## Declaration of Generative AI and AI-assisted technologies in the writing process

During the preparation of this work the authors used ChatGPT in order to assist in phrasing certain sentences. After using this tool/service, the authors reviewed and edited the content as needed and take full responsibility for the content of the publication.

## References

Bengel, P., Dybkova, N., Tirilomis, P., Ahmad, S., Hartmann, N. B A. M., Krekeler, M. C., Maurer, W., Pabel, S., Trum, M., Mustroph, J., Gummert, J., Milting, H., Wagner, S., Ljubojevic-Holzer, S., Toischer, K., Maier, L. S., Hasenfuss, G., Streckfuss-Bomeke, K., & Sossalla, S. (2021). Detrimental proarrhythmogenic interaction of Ca(2+)/calmodulin-dependent protein kinase II and Na(V)1.8 in heart failure. Nat Commun, 12(1), 6586. 10.1038/s41467-021-26690-1

Borchert, T., Hubscher, D., Guessoum, C. I., Lam, T. D., Ghadri, J. R., Schellinger, I. N., Tiburcy, M., Liaw, N. Y., Li, Y., Haas, J., Sossalla, S., Huber, M. A., Cyganek, L., Jacobshagen, C., Dressel, R., Raaz, U., Nikolaev, V. O., Guan, K., Thiele, H., … Streckfuss-Bomeke, K. (2017). Catecholamine-Dependent beta-Adrenergic Signaling in a Pluripotent Stem Cell Model of Takotsubo Cardiomyopathy. J Am Coll Cardiol, 70(8), 975–991. 10.1016/j.jacc.2017.06.061

Cyganek, L., Tiburcy, M., Sekeres, K., Gerstenberg, K., Bohnenberger, H., Lenz, C., Henze, S., Stauske, M., Salinas, G., Zimmermann, W. H., Hasenfuss, G., & Guan, K. (2018). Deep phenotyping of human induced pluripotent stem cell-derived atrial and ventricular cardiomyocytes. JCI Insight, 3(12). 10.1172/jci.insight.99941

Dridi, H., Kushnir, A., Zalk, R., Yuan, Q., Melville, Z., & Marks, A. R. (2020). Intracellular calcium leak in heart failure and atrial fibrillation: a unifying mechanism and therapeutic target. Nat Rev Cardiol, 17(11), 732–747. 10.1038/s41569-020-0394-8

Ebrahimi, Z., Loni, M., Daneshtalab, M., & Gharehbaghi, A. (2020). A review on deep learning methods for ECG arrhythmia classification. Expert Systems with Applications: X, 7, 100033. 10.1016/j.eswax.2020.100033

El-Battrawy, I., Cammann, V. L., Kato, K., Szawan, K. A., Di Vece, D., Rossi, A., Wischnewsky, M., Hermes-Laufer, J., Gili, S., Citro, R., Bossone, E., Neuhaus, M., Franke, J., Meder, B., Jaguszewski, M., Noutsias, M., Knorr, M., Heiner, S., D’Ascenzo, F., … Templin, C. (2021). Impact of Atrial Fibrillation on Outcome in Takotsubo Syndrome: Data From the International Takotsubo Registry. J Am Heart Assoc, 10(15), e014059. 10.1161/JAHA.119.014059

El-Battrawy, I., Lang, S., Ansari, U., Behnes, M., Hillenbrand, D., Schramm, K., Fastner, C., Zhou, X., Bill, V., Hoffmann, U., Papavassiliu, T., Elmas, E., Haghi, D., Borggrefe, M., & Akin, I. (2017). Impact of concomitant atrial fibrillation on the prognosis of Takotsubo cardiomyopathy. Europace, 19(8), 1288–1292. 10.1093/europace/euw293

G Van Rossum, F. D. (1995). Python reference manual. Corporation for National Research Initiatives (CNRI) in Reston, Virginia

Guatimosim, S., Guatimosim, C., & Song, L. S. (2011). Imaging calcium sparks in cardiac myocytes. Methods Mol Biol, 689, 205–214. 10.1007/978-1-60761-950-5_12

Hartmann, N., Knierim, M., Maurer, W., Dybkova, N., Hasenfuss, G., Sossalla, S., & Streckfuss-Bomeke, K. (2023). Molecular and Functional Relevance of Na(V)1.8-Induced Atrial Arrhythmogenic Triggers in a Human SCN10A Knock-Out Stem Cell Model. Int J Mol Sci, 24(12). 10.3390/ijms241210189

Hartmann, N., Knierim, M., Maurer, W., Dybkova, N., Zeman, F., Hasenfuß, G., Sossalla, S., & Streckfuss-Bömeke, K. (2024). Na(V)1.8 as Proarrhythmic Target in a Ventricular Cardiac Stem Cell Model. Int J Mol Sci, 25(11). 10.3390/ijms25116144

Jordà, P., Toro, R., Diez, C., Salazar-Mendiguchía, J., Fernandez-Falgueras, A., Perez-Serra, A., Coll, M., Puigmulé, M., Arbelo, E., García-Álvarez, A., Sarquella-Brugada, G., Cesar, S., Tiron, C., Iglesias, A., Brugada, J., Brugada, R., & Campuzano, O. (2021). Malignant Arrhythmogenic Role Associated with RBM20: A Comprehensive Interpretation Focused on a Personalized Approach. J Pers Med, 11(2). 10.3390/jpm11020130

Karakikes, I., Ameen, M., Termglinchan, V., & Wu, J. C. (2015). Human induced pluripotent stem cell-derived cardiomyocytes: insights into molecular, cellular, and functional phenotypes. Circ Res, 117(1), 80–88. 10.1161/CIRCRESAHA.117.305365

Lian, X., Zhang, J., Azarin, S. M., Zhu, K., Hazeltine, L. B., Bao, X., Hsiao, C., Kamp, T. J., & Palecek, S. P. (2013). Directed cardiomyocyte differentiation from human pluripotent stem cells by modulating Wnt/beta-catenin signaling under fully defined conditions. Nat Protoc, 8(1), 162–175. 10.1038/nprot.2012.150

Masarone, D., Limongelli, G., Rubino, M., Valente, F., Vastarella, R., Ammendola, E., Gravino, R., Verrengia, M., Salerno, G., & Pacileo, G. (2017). Management of Arrhythmias in Heart Failure. J Cardiovasc Dev Dis, 4(1). 10.3390/jcdd4010003

Motiejunaite, J., Amar, L., & Vidal-Petiot, E. (2021). Adrenergic receptors and cardiovascular effects of catecholamines. Ann Endocrinol (Paris), 82(3-4), 193–197. 10.1016/j.ando.2020.03.012

Novak, A., Barad, L., Lorber, A., Gherghiceanu, M., Reiter, I., Eisen, B., Eldor, L., Itskovitz-Eldor, J., Eldar, M., Arad, M., & Binah, O. (2015). Functional abnormalities in iPSC-derived cardiomyocytes generated from CPVT1 and CPVT2 patients carrying ryanodine or calsequestrin mutations. J Cell Mol Med, 19(8), 2006–2018. 10.1111/jcmm.12581

Oguntuyo, K., Schuftan, D., Guo, J., Simmons, D., Bhagavan, D., Moreno, J. D., Kang, P. W., Miller, E., Silva, J. R., & Huebsch, N. (2022). Robust, Automated Analysis of Electrophysiology in Induced Pluripotent Stem Cell-Derived Micro-Heart Muscle for Drug Toxicity. Tissue Eng Part C Methods, 28(9), 457–468. 10.1089/ten.tec.2022.0053

Pabel, S., Knierim, M., Stehle, T., Alebrand, F., Paulus, M., Sieme, M., Herwig, M., Barsch, F., Kortl, T., Poppl, A., Wenner, B., Ljubojevic-Holzer, S., Molina, C. E., Dybkova, N., Camboni, D., Fischer, T. H., Sedej, S., Scherr, D., Schmid, C., … Sossalla, S. (2022). Effects of Atrial Fibrillation on the Human Ventricle. Circ Res, 130(7), 994–1010. 10.1161/CIRCRESAHA.121.319718

Psaras, Y., Margara, F., Cicconet, M., Sparrow, A. J., Repetti, G. G., Schmid, M., Steeples, V., Wilcox, J. A. L., Bueno-Orovio, A., Redwood, C. S., Watkins, H. C., Robinson, P., Rodriguez, B., Seidman, J. G., Seidman, C. E., & Toepfer, C. N. (2021). CalTrack: High-Throughput Automated Calcium Transient Analysis in Cardiomyocytes. Circ Res, 129(2), 326–341. 10.1161/CIRCRESAHA.121.318868

Rebs, S., Sedaghat-Hamedani, F., Kayvanpour, E., Meder, B., & Streckfuss-Bomeke, K. (2020). Generation of pluripotent stem cell lines and CRISPR/Cas9 modified isogenic controls from a patient with dilated cardiomyopathy harboring a RBM20 p.R634W mutation. Stem Cell Res, 47, 101901. 10.1016/j.scr.2020.101901

Sammani, A., Kayvanpour, E., Bosman, L. P., Sedaghat-Hamedani, F., Proctor, T., Gi, W. T., Broezel, A., Jensen, K., Katus, H. A., Te Riele, A., Meder, B., & Asselbergs, F. W. (2020). Predicting sustained ventricular arrhythmias in dilated cardiomyopathy: a meta-analysis and systematic review. ESC Heart Fail, 7(4), 1430–1441. 10.1002/ehf2.12689

Shah, M., Akar, F. G., & Tomaselli, G. F. (2005). Molecular basis of arrhythmias. Circulation, 112(16), 2517–2529. 10.1161/CIRCULATIONAHA.104.494476

Ter Keurs, H. E., & Boyden, P. A. (2007). Calcium and arrhythmogenesis. Physiol Rev, 87(2), 457–506. 10.1152/physrev.00011.2006

Tohyama, S., Hattori, F., Sano, M., Hishiki, T., Nagahata, Y., Matsuura, T., Hashimoto, H., Suzuki, T., Yamashita, H., Satoh, Y., Egashira, T., Seki, T., Muraoka, N., Yamakawa, H., Ohgino, Y., Tanaka, T., Yoichi, M., Yuasa, S., Murata, M., Fukuda, K. (2013). Distinct metabolic flow enables large-scale purification of mouse and human pluripotent stem cell-derived cardiomyocytes. Cell Stem Cell, 12(1), 127–137. 10.1016/j.stem.2012.09.013

