## Supplementary Information for "Arrhythpy: An Automated Tool to Quantify and Classify Arrhythmias in Ca^2+^ Transients of iPSC-Cardiomyocytes"

**Supplements:**

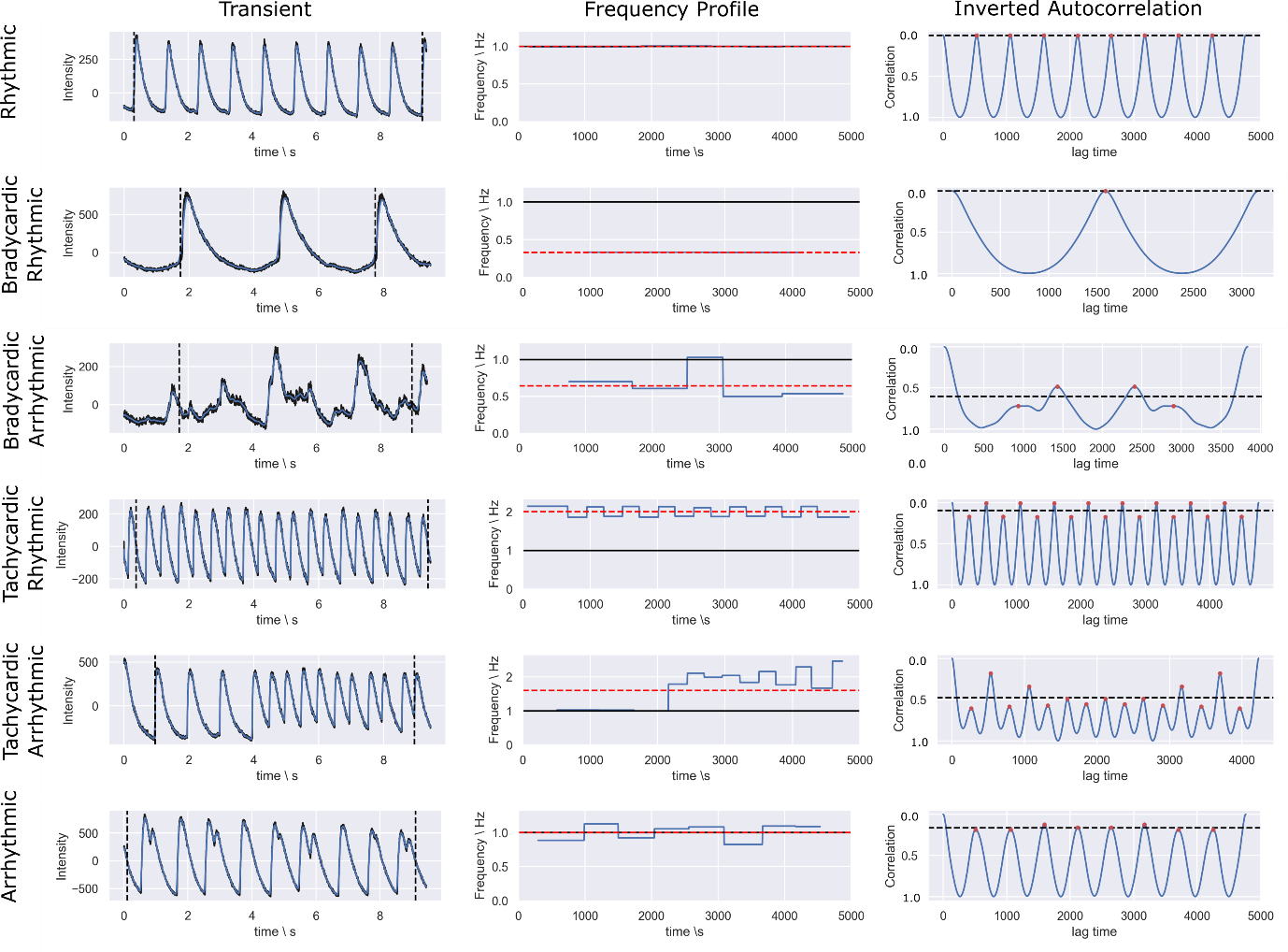

Fig. S1: Example transients (left column) and the subsequent analysis consisting of wavelet transformation obtaining the frequency profile (middle column) and the inverse autocorrelation (right column).

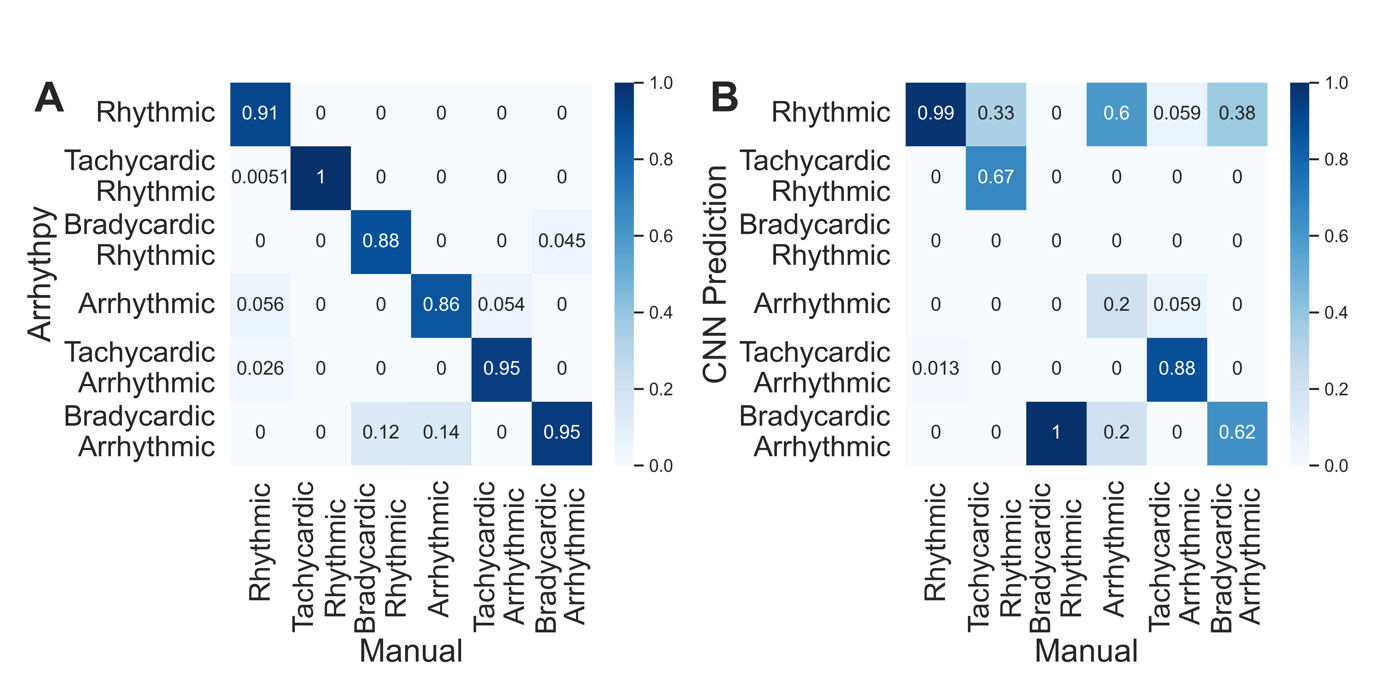

Fig. S2: Comparison of *Arrhthpy-based* classification and CNN-based classification. A) Confusion matrix shows high values along the diagonal, indicating good classification performance of Arrhythpy. B) Confusion matrix of the CNN classification shows that the CNN is very accurate for rhythmic transients but fails to classify the arrhythmias correctly due to the lack of labeled data.

|  | Mean Frequency | Frequency Variance | Bradycardia | Tachycardia | Mean Peak Correlation | Measure of Arrhythmia |
| --- | --- | --- | --- | --- | --- | --- |
| Rhythmic | 1.00 | 0.00 | 0.00 | 0.00 | 0.00 | 0.00 |
| Bradycardic Rhythmic | 0.33 | 0.00 | 0.67 | 0.00 | 0.00 | 0.00 |
| Bradycardic Arrhythmic | 0.65 | 0.03 | 0.39 | 0.00 | 0.60 | 0.31 |
| Tachycardic Rhythmic | 2.01 | 0.02 | 0.00 | 1.01 | 0.09 | 0.06 |
| Tachycardic  Arrhythmic | 1.60 | 0.25 | 0.00 | 0.60 | 0.48 | 0.36 |
| Arrhythmic | 0.99 | 0.11 | 0.05 | 0.05 | 0.16 | 0.14 |

Table S1: Parameters obtained by analyzing the transients in Fig. S1 with *Arrhythpy.* The classification is based on setting the threshold on Bradycardia/Tachycardia to 0.1 and the threshold on the measure of arrhythmia to 0.1.

| **Fig. 4:**  **DCM**  **ventricle** | **Basal** | **Iso (1µM)** |  |
| --- | --- | --- | --- |
|  | 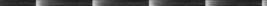  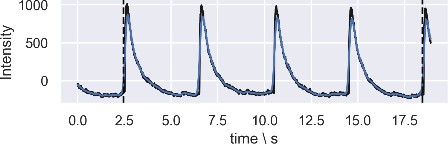 | 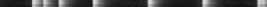  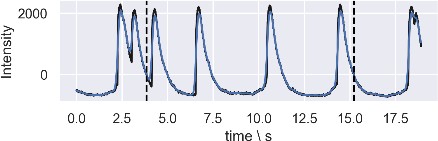 |  |
| **Fig. 5:**  **CTRL**  **atrium** | **Basal** | **Iso (50nM)** |  |
|  | 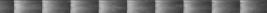  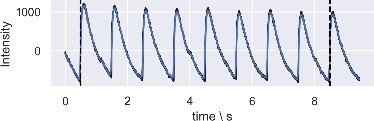 | 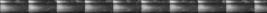  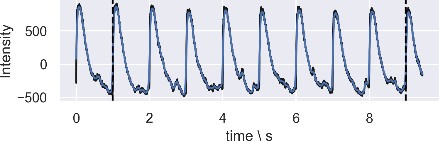 |  |
| **Fig. 5:**  **TTS**  **atrium** | **Basal** | **Iso (50 nM)** | **Metoprolol** |
|  | 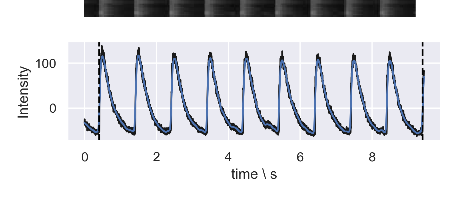 | 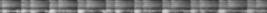  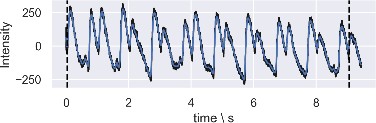 | 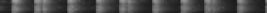  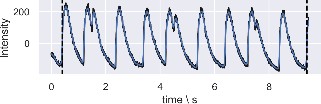 |

Table S2: Examplary line scans and respective transients of a ventricular DCM and atrial CTRL cell (basal and Iso treatment) and an atrial TTS cell (basal, Iso and Metoprolol treatment).
